# Clinical long-read sequencing of the human mitochondrial genome for mitochondrial disease diagnostics

**DOI:** 10.1101/597187

**Authors:** Elizabeth Wood, Matthew D Parker, Mark J Dunning, Sirisha Hesketh, Dennis Wang, Ryan Pink, Carl Fratter

## Abstract

**Purpose:** Long-read, third generation, sequencing technologies have the potential to improve current state of the art diagnostic strategies. In order to determine if long-read sequencing technologies are suitable for the diagnosis of mitochondrial disorders due to mitochondrial DNA (mtDNA) variants, particularly large deletions, we compared the performance of Oxford Nanopore Technologies (ONT) MinION to current diagnostic methods.

**Methods:** We sequenced mtDNA from nine patients with mtDNA deletion disorders and three normal controls with both ONT MinION and Illumina MiSeq. We applied a computational pipeline to estimate the positions of mtDNA deletions in patients, and subsequently validated the breakpoints using Sanger sequencing.

**Results:** We were able to detect mtDNA deletions with a MinION workflow, successfully calling the disease causing event in all cases. Sequencing coverage was in most cases significantly more (p=0.03, Wilcoxon test) uniform with MinION than with MiSeq and subsequent correction of MinION reads improved breakpoint accuracy and reduced false positives. Although heteroplasmic single nucleotide variants are detectable, the high number of false positives and false negatives precludes their use in diagnostics at this time.

**Conclusion:** The MinION is becoming an increasingly attractive diagnostic tool due to the reducing cost, increasing accuracy, and the speed at which data can be obtained.

## Introduction

Mitochondria are organelles that are responsible for energy production via oxidative phosphorylation, which takes place in the mitochondrial respiratory chain. Mitochondrial DNA (mtDNA) is a 16.6kb closed-circle present in hundreds to thousands of copies per cell. This DNA encodes thirteen subunits of the respiratory chain.

Mitochondrial diseases are characterised by biochemical abnormalities of the respiratory chain. Genetic causes include pathogenic mtDNA aberrations (primary mtDNA disorders) and pathogenic aberrations in nuclear genes that affect mtDNA maintenance. The majority of pathogenic mtDNA variants are heteroplasmic (a mixture of wild-type and mutant mtDNA) with a threshold level of heteroplasmy above which mitochondrial function is impaired^1^.

Here we focus on an important subset of primary mtDNA disorders caused by large (2-8kb), usually sporadic, heteroplasmic mtDNA deletions.

Current routine diagnostic testing strategies for mtDNA deletions include polymerase chain reaction (PCR) and Southern blotting. Long-range PCR (LR-PCR) is used to amplify the mitochondrial genome, followed by agarose gel electrophoresis. Any fragments smaller than the expected size are suggestive of deletions; however this low-resolution approach doesn’t provide the location, or indeed a truly accurate deletion size. Southern blotting can be used to determine the approximate deletion size and estimate the level of heteroplasmy. However, it is labour-intensive, time-consuming, and cannot determine the precise deletion location.

The rapid developments in 3rd generation sequencing technologies make them attractive methods for use in clinical diagnostics. Pacific Biosciences (PacBio) and Oxford Nanopore Technologies (ONT) have recently developed sequencing technologies that allow DNA fragments tens of kilobases in length to be sequenced. This introduces the possibility of sequencing the full mitochondrial genome in a single read.

To improve the clinical characterisation of mtDNA deletions and determine the feasibility of using ONT MinION long-read sequencing in a clinical diagnostic pathway, we used LR-PCR to amplify the mitochondrial genome from patients with known aberrations (and controls) followed by both Illumina MiSeq and MinION sequencing.

## Materials & Methods

### Sample Selection

Fifteen patients were selected based on previous mitochondrial investigations; nine with single deletions, two with multiple deletions, and four negative controls (Table S1).

### Long Range PCR

Primers were designed to amplify 16.1kb of the mitochondrial genome and LA Taq Hot Start polymerase (TaKaRa Bio, Japan) was used to generate LR-PCR products. The products were quantified and each sample divided for MiSeq and MinION library preparation.

### MinION Library Preparation and Sequencing

A MinION library was produced using the ONT Ligation Sequencing Kit 1D (SQK-LSK108) and Native Barcoding Kit 1D (EXP-NBD103), according to the 1D Native Barcoding Genomic DNA Protocol. 1 μg LR-PCR DNA was used for End-repair/dA-tailing, followed by barcoding. Equimolar amounts of barcoded samples were pooled. Adapters were ligated and loaded onto an R9 flow cell (FLO-MIN106) and sequenced for 40hrs.

### MinION Data Analysis

Fast5 files were converted into fastq files with poretools (https://www.biorxiv.org/content/early/2014/07/23/007401 - Unpublished v0.6.0). Adapters were trimmed and reads demultiplexed with porechop (Unpublished v0.2.3). Reads were either mapped to the revised Cambridge reference sequence (rCRS; NC_012920.1) using minimap2^2^ (v2.13) or were corrected and trimmed with canu^3^ (v1.7.1) before mapping. Alignments were sorted and coverage calculated with samtools^4^ (v1.9). Structural Variants (SV)s were called with NanoSV^5^ (v1.2.2). Small variants were detected with VarScan^6^ (v2.3.9) after re-calling bases with albacore (v2.3.3 - Oxford Nanopore). Code for this pipeline can be found at https://github.com/sheffield-bioinformatics-core/mMinION. This pipeline is summarised in Figure S5.

### MiSeq Library Preparation and Sequencing

The NexteraXT DNA Sample Preparation Kit (Illumina) was used to make barcoded libraries from the LR-PCR products and subsequently sequenced on the Illumina MiSeq.

### MiSeq Data Analysis

Reads were mapped to rCRS with bwa^7^ mem (v0.7.17), duplicates were marked with Picard MarkDuplicates (v2.18.11, https://github.com/broadinstitute/picard). Coverage was calculated using samtools depth with a minimum quality of 20. Read groups were added with Picard. SVs were detected using LUMPY v0.2.13^8^ and genotyped with svtyper v0.7.0^9^ Sequencing statistics can be found in Figure S3.

### Breakpoint Validation

Primer sets (details on request) were used to amplify across the putative breakpoints. Sanger sequencing was carried out on an ABI-3730 (Thermo Fisher Scientific).

Further details are described in Supplementary Methods.

## Results

### PCR of Mitochondrial Genome

Figure S1 shows the products of LR-PCR for the 15 samples in this study. Based on these results and to be amenable to a 12 samples barcoded library, 12 patients were chosen for sequencing, as indicated in Table S1.

#### Increased Uniformity of Coverage Using Oxford Nanopore Sequencing

A single multiplexed MinION run was performed producing 150,607 reads marked as pass by the MinION totaling 0.750 gigabases. Reads spanning the entire amplicon were generated, but the median read length was 3.2kb (Figure S2). Reads longer than the amplicon length in each case were the result of chimeric fragments containing more than one copy of mtDNA (Figure S4).

Mapping MinION reads to rCRS with minimap2 resulted in alignments that exhibit reductions in coverage corresponding to the deletions present in all positive cases (Figure 1C & S7). Due to the heteroplasmic nature of the mtDNA deletions, wild-type mitochondrial genomes were detected at a very low frequency, likely due to the preferential amplification of the shorter mtDNA fragments containing deletions.

**Figure 1.**
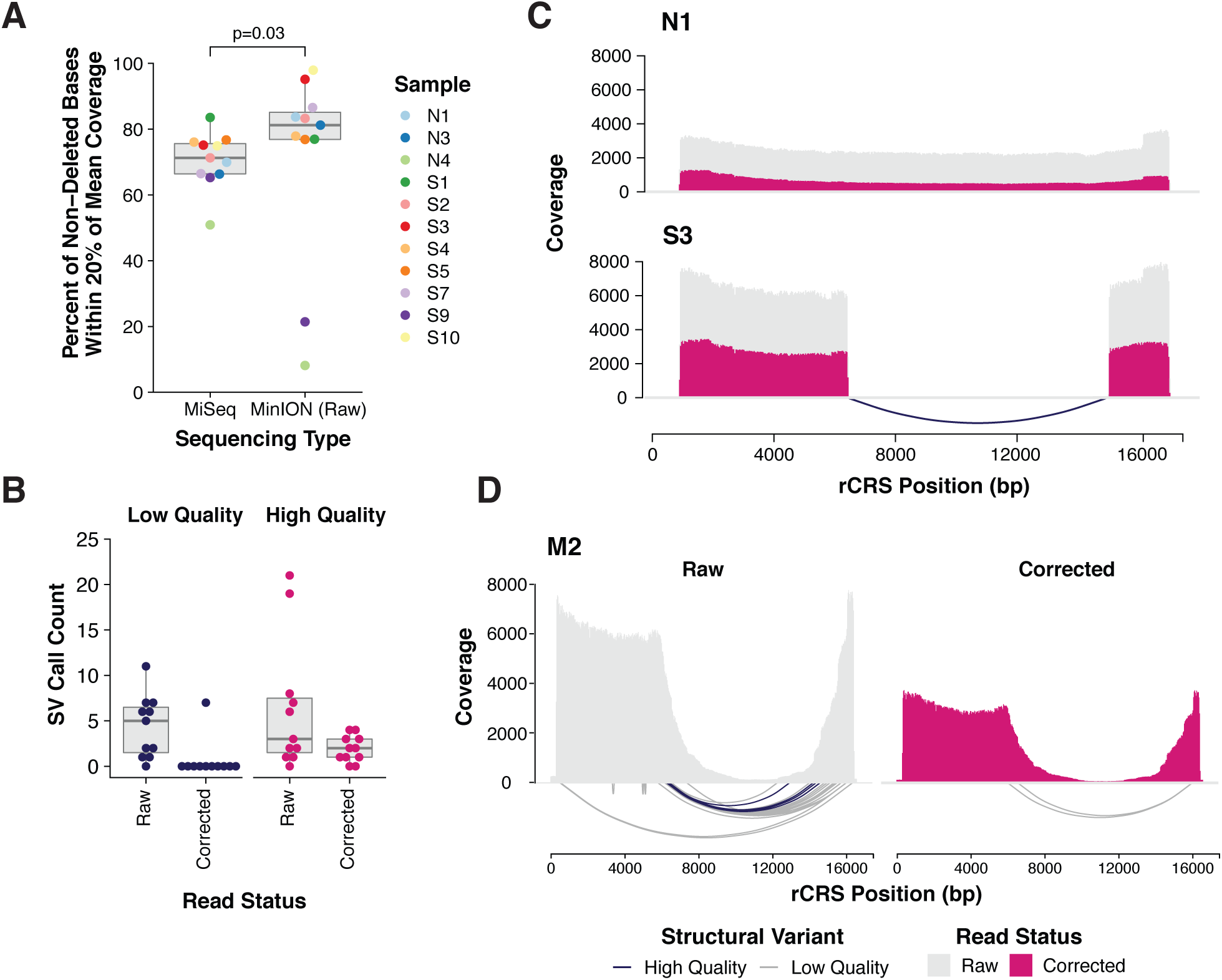
MinION Sequencing of the Mitochondrial Genome. **A.** Coverage uniformity was calculated in all samples for both MiSeq and MinION generated data. MinION produced a statistically significant (p=0.03) increase in uniformity (Wilcoxon test). **B.** After canu correction the total number of structural variant (SV) predictions using NanoSV is decreased and the proportion of these which are high quality is increased. **C.** Coverage along the mitochondrial genome showing both raw (grey) and canu corrected & trimmed reads (pink) for samples N1 and S3. Deletions are represented by defined drops in coverage in all patients with mtDNA deletions. SV predictions are represented by lines under the histogram coloured by their quality. SVs resulting from artefacts at the start and end (< 320 and > 16400) of the amplicon and those SVs labelled as INS by NanoSV were not plotted. **D.** Coverage and SV predictions for sample M2, containing multiple deletions, for both raw and corrected reads.

Interestingly, coverage uniformity is increased (p=0.03, Wilcoxon test) when using MinION to sequence the mitochondrial amplicons when compared to the MiSeq (Figure 1A).

#### Computational Structural Variant Detection

To determine if we could use Structural Variant (SV) calling to predict the exact location of deletions, we applied NanoSV to two sets of mapped reads; 1) those originating from raw reads, and 2) canu corrected and trimmed reads. Correction and trimming made a marked difference to the quality of MinION reads, creating a consensus that helped remove random error (Figure S6). Recent benchmarking of correction tools suggested that canu outperforms most tools for this purpose (https://www.biorxiv.org/content/10.1101/519330v1 - Unpublished).

Using raw reads and NanoSV we were able to detect SVs responsible for the deletion in all single deletion samples (Table S3), although the total number of SV calls was relatively high, and N1, N3 and N4 each contained a high quality SV call. Correction and trimming of reads using canu (Figure S6) improved NanoSV predictions; reducing the number of false positives in each case (Tables S2 & S3) and resulting in refinement of the breakpoint positions. Moreover, high quality calls were now absent from N1 and N3 (Figure S7 & Table S2). In all samples, except S7, the exact breakpoint position was not be determined presumably due to homology around the 5’ and the 3’ ends of the deletion breakpoints (Figure S9).

Sample M2 is predicted to contain multiple deletions at varying levels of heteroplasmy. From the coverage plots (Figure 1D), it is clear that this sample has a complex collection of deletions. Computational SV callers can help elucidate the exact configuration of SVs. However, as this sample possesses multiple heteroplasmic deletion events it was deemed not suitable for correction and trimming. Methods such as canu attempt to find a consensus sequence, which would result in a loss of evidence for events with low levels of supporting reads (Figure 1D - Right panel).

Despite Sample N4 showing coverage changes that could be mistaken for deletions in this patient (Figure S7), SV calling resulted in the detection of only one high quality structural variant out of a total of 8. Since no high quality predictions were present in the MiSeq data for this sample (Table S4), the coverage fluctuations are therefore likely an artefact of PCR and subsequent sequencing.

Interestingly the total number of SVs predicted using LUMPY express and svtyper on MiSeq data was high compared to MinION, although the number of high quality variants was a small fraction of this total (Table S4). No normal samples had high quality SVs, however with default parameters we also failed to detect any high quality SVs acceptably close to the Sanger confirmed breakpoints in MiSeq data from samples S3 and S9.

Sanger sequencing was carried out across the breakpoints for the single deletion patients (Figure S9). In sample S7 this confirmed the breakpoints determined by NanoSV as m.7642, located within the *MT-CO2* gene, and m.12985, located within the *MT-ND5* gene. Sanger results for the remaining single deletion patients confirmed the presence of the predicted deletion and a short flanking region of homology.

#### Small Variant Detection

After re-calling bases with albacore, we detected single nucleotide variants (SNV) with VarScan in both the MinION and MiSeq data. Samples N1 and N4 contained previously known heteroplasmic pathogenic variants at a frequency of 22% (m.3243A>G) and 89% (m.14430A>G) respectively. Figures S11 and S12 show allele frequency comparison plots and summaries of the number of variants called by each technology. Increasing the minimum average base quality (min-avg-qual) required to support a variant from 15 to 20 decreased the total number of variants called. For example, in sample N1, 401 variants were called with a min-avg-qual of 15; only 33 of these were shared between MiSeq and MinION, and 1 was found in MiSeq data only. This suggests a high false positive rate when using MinION data. Increasing the min-avg-qual to 20, however, resulted in 59 total calls, 32 of these were shared, and 2 in MiSeq only. This trend was mirrored in other samples (Figure S12).

To assess the MinION’s ability to detect and quantify heteroplasmic variants, we investigated the variant calls for the heteroplasmic pathogenic variants in samples N1 and N4 (m.3243A>G and m.14430A>G respectively). Applying minimum base quality cut-offs of 15 and 20 resulted in allele frequencies of 23.86% and 28.52% of the variant in sample N1 on the MinION. On the MiSeq, the variant showed the same allele frequency of 23.18% when the cut-off was increased from 15 to 20. However, in sample N4, setting a higher quality threshold dramatically reduced the allele frequency, from 58.46% to 23.68% whereas at both quality thresholds the frequency was 89.49% in data generated by the MiSeq. Although these results are promising, further work is required to refine small variant calling in MinION data. Unlike for SV calling, we were unable to use read correction and trimming for this application because low frequency variants are lost from the data (Figure S10).

## Discussion

We have shown that long-read sequencing is an emerging strategy to characterise the breakpoints of major mtDNA deletions. Single and multiple mtDNA deletions were detected and characterised in both MinION and MiSeq data.

A major advantage of using MinION over MiSeq was a significantly increased uniformity of coverage of the mtDNA amplicon. Additionally the MinION was able to produce reads covering the full 16.1kb amplicon. However, many shorter reads (in total approximately 17% of reads were below 1kb in length) were also present mainly at the start and end of the amplicon, sometimes resulting in false positive SV calls. It is likely that the shorter fragments were generated by shearing during the library preparation, and filtering these bioinformatically introduces false negatives if structural variants are present close to the beginning or end of the amplicon. Further experimental optimisation is required to reduce the production of these fragments.

Using NanoSV we were able to detect SVs explaining the deletions in all patients in MinION data. Due to homology at deletion breakpoints (a previously reported feature of mtDNA deletions^10–12^), which ranged from 8-13 nucleotides, NanoSV was unable to call the exact bases at which the deletion had occurred. Canu correction and trimming improved structural variant accuracy and in addition dramatically reduced false positive calls. Correction, however, is not suitable when low frequency heteroplasmic events are present because, when finding a consensus, low frequency events are excluded from the final corrected reads. SV calling in the MinION data was superior to that for the MiSeq data (using LUMPY Express, and svtyper), which was unable to find deletion breakpoints in S3 and S9. All breakpoints in single deletion patients were confirmed with Sanger sequencing.

SNV calling in MinION data was also evaluated. We were able to detect known heteroplasmic pathogenic SNVs in samples N1 and N4. In general there was surprisingly good concordance with variants called using MiSeq data although the number of false positive calls was high.

Both the short-read (Illumina MiSeq) and long-read (ONT MinION) strategies allow the elucidation of breakpoint locations in patients with mtDNA deletions, helping to facilitate more detailed genotype/phenotype correlations for these mitochondrial disorders. These next generation sequencing strategies are therefore superior to current routine practice of LR-PCR and/or Southern blotting. In addition, the simultaneous detection of deletions and SNVs provides a single laboratory strategy for all primary mtDNA disorders, as previously described for short-read NGS^13^. Due to the high error rate of long-read sequencing, this technology is not currently appropriate for the detection of SNVs as well as large deletions. However, owing to the shorter library preparation and sequencing time, the generation of single reads covering the entire mitochondrial genome, and uniform coverage, ONT sequencing has potential for use in mtDNA diagnostics if sequencing accuracy continues to improve as anticipated. In addition, ONT MinION’s low capital costs may allow the introduction of mtDNA NGS into diagnostic centres where costs are prohibitive.

## Acknowledgements

This research was supported / funded by the NIHR Sheffield Biomedical Research Centre (BRC) / NIHR Sheffield Clinical Research Facility (CRF), the UK NHS Scientist Training Programme, the Oxford Centre of the Rare Mitochondrial Disorders Service (mitochondrialdisease.nhs.uk, funded by UK NHS Highly Specialised Services), and Oxford Brookes University and The Royal Society. The views expressed are those of the author(s) and not necessarily those of the NHS, the NIHR or the Department of Health and Social Care (DHSC). The authors would like to thank Natalie Groves for useful discussions around the analysis of MinION Data. The authors would also like to thank all other staff in Oxford Medical Genetics Laboratories who supported this project, in particular Philip Hodsdon for technical expertise and support.

