## Supplementary Information for "Clinical long-read sequencing of the human mitochondrial genome for mitochondrial disease diagnostics"

#### Table of Contents

**Supplementary Methods** **22**
Long-range PCR 22
MiSeq Library Preparation and Sequencing 23
Coverage Uniformity Calculation 23
**Supplementary Tables** **24**
Table S1. Samples Selected for Sequencing 24
Table S2- Sanger Sequencing Results & MinION Structural Variant Prediction using NanoSV after Canu Correction & Trimming 25
Table S3 - MinION Raw Structural Variant Prediction 26
Table S4 - MiSeq Structural Variant Prediction 27
**Supplementary Figures** **28**
Figure S1 - PCR of Mitochondrial DNA 28
Figure S2 - MinION Sequencing Statistics 28
Figure S3 - MiSeq Sequencing Statistics 29
Figure S4 - Read Lengths in Sample S10 30
Figure S5 - MinION SV analysis pipeline 31
Figure S6 - Canu Correction of MinION Reads 32
Figure S7 - MinION Coverage and High Quality SV Predictions 33
Figure S8 - MiSeq Coverage and High Quality SV Predictions 34
Figure S9 - Sequence Homology Around Deletion Breakpoints 35
Figure S10 - Correction causes loss of known SNV 37
Figure S11 - Known variants are called by VarScan using MinION data 38
Figure S12 - Variant Calling Comparison for All Samples 39
**Supplementary References** 40

### Supplementary Methods

#### Long-range PCR

Primers were designed to amplify 16.1kb of the mitochondrial genome (forward primer 5'-CCGCTTCTGGCCACAGCACTTAAACAC-3', reverse primer 5'-GGAGGATGGTGGTCAAGGGACCCCTAT-3') and the LA Taq Hot Start Version polymerase (TaKaRa Bio, Japan) was used to generate long range PCR products. Reactions were carried out in a volume of 20 μl using ~35 ng template DNA, with 1 unit polymerase, 0.5 μM primers and 0.4 mM dNTPs. The PCR conditions were: 94°C 1 minute, 30 cycles [98°C 10 seconds, 68°C 12 minutes], 72°C 10 minutes. Twelve of the 15 amplicons were selected for sequence analysis. The products were quantified using a GloMax Multi+ Fluorometer and each sample was divided for MiSeq and MinION library preparation.

#### MiSeq Library Preparation and Sequencing

The concentration of each LR-PCR product was reduced to 0.2 ng/μl by three sequential rounds of quantification and dilution. A sequencing library was then made using the NexteraXT DNA Sample Preparation Kit (Illumina). An engineered transposome was used to fragment the DNA (to fragments of ~300bp) alongside adapter ligation; a limited cycle PCR reaction (12 cycles) was then used to add dual-indexing tags and library quality control was carried out using the Agilent TapeStation. Equimolar amounts of each barcoded sample were then pooled for sequencing on the Illumina MiSeq.

#### Coverage Uniformity Calculation

Coverage uniformity was defined as the percent of bases with coverage within 20% of the total mean coverage for the amplicon excluding known deleted bases and positions <320 and >16400. Only bases with a quality of >= 20 were included in the calculation. Statistical significance was determined with a Wilcoxon test.

### Supplementary Tables

| Sample ID | Sample Type | Deletion Size (kb)* | Sequenced | Notes |
|---|---|---|---|---|
| **Single Deletion** | | | | |
| S1 | Muscle | 5.0 | Yes | - |
| S2 | Blood | 5.0 | Yes | - |
| S3 | Muscle | 9.4 | Yes | - |
| S4 | Blood | 5.0 | Yes | - |
| S5 | Bone Marrow | 4.4 | Yes | - |
| S6 | Muscle | 7.4 | No | - |
| S7 | Muscle | 5.6 | Yes | - |
| S9 | Muscle | 2.0 | Yes | - |
| S10 | Fibroblast | 7.8 | Yes | - |
| **Multiple Deletions** | | | | |
| M2 | Muscle | NA | Yes | - |
| M3 | Muscle | NA | No | - |
| **Normal Controls** | | | | |
| N1 | Blood | NA | Yes | 22% m.3243A>G |
| N2 | Blood | NA | No | 11% m.3243A>G |
| N3 | Muscle | NA | Yes | - |
| N4 | Muscle | NA | Yes | 89% m.14430A>G |

**Table S1. Samples Selected for Sequencing**

*Deletion sizes are estimated from Southern blot analysis.

| | Southern | Sanger | | | | MinION | | | |
|---|---|---|---|---|---|---|---|---|---|
| | Size (kb) | Start* | End* | Size (kb) | HGVS | Start | End | QUAL* | HQ SVs |
| **Normal Controls** | | | | | | | | | |
| N1 | - | - | - | | | - | - | - | 0 |
| N3 | - | - | - | | | - | - | - | 0 |
| N4 | - | - | - | | | - | - | - | 1 (8) |
| **Single Deletion Samples** | | | | | | | | | |
| S1 | 5.0 | 9525 | 14387 | 4.86 | m.9525_14387del | 9529 | 14377 | 9999 | 1 (1) |
| S2 | 5.0 | 8483 | 13459 | 4.98 | m.8483_13459del | 8469 | 13447 | 9999 | 4 (4) |
| | | | | | | 8446 | 13485 | 53 | |
| S3 | 9.4 | 5850 | 14423 | 8.57 | m.5850_14423del | 5846 | 14419 | 9999 | 2 (2) |
| S4 | 5.0 | 8483 | 13459 | 4.98 | m.8483_13459del | 8469 | 13447 | 9999 | 1 (1) |
| S5 | 4.4 | 9498 | 13734 | 4.24 | m.9498_13734del | 9487 | 13726 | 9999 | 3 (3) |
| S7 | 5.6 | 7642 | 12984 | 5.34 | m.7642_12984del | 7641 | 12985 | 9999 | 3 (3) |
| S9 | 2.0 | 12113 | 14421 | 2.31 | m.12113_14421del | 12108 | 14412 | 9999 | 2 (2) |
| S10 | 7.8 | 8649 | 16084 | 7.44 | m.8649_16084del | 8648 | 16073 | 9999 | 4 (4) |

* QUAL - Quality score

**Table S2- Sanger Sequencing Results & MinION Structural Variant Prediction using NanoSV after Canu Correction & Trimming**

* Where repeat regions are present at the breakpoints, the most 3' position is given.

| | Sanger | | MinION | | | |
|---|---|---|---|---|---|---|
| **Sample** | **Start** | **End** | **Start** | **End** | **QUAL** | **Total SVs** |
| N1 | - | - | - | - | - | 1 (3) |
| N3 | - | - | - | - | - | 0 (1) |
| N4 | - | - | - | - | - | 1 (8) |
| S1 | 9525 | 14387 | 9529 | 14377 | 9999 | 2 (2) |
| S2 | 8483 | 13459 | 8473 | 13449 | 9999 | 19 (26) |
| | | | 8483 | 8431 | 2725 | |
| S3 | 5850 | 14423 | 5846 | 14419 | . | 3 (9) |
| | | | 5831 | 14506 | 157 | |
| S4 | 8483 | 13459 | 8473 | 13448 | 9999 | 6 (8) |
| S5 | 9498 | 13734 | 9488 | 13724 | 9999 | 21 (26) |
| S7 | 7642 | 12984 | 7641 | 12985 | . | 8 (14) |
| | | | 7618 | 13059 | 203 | |
| S9 | 12113 | 14421 | 12107 | 14412 | 9999 | 2 (3) |
| S10 | 8649 | 16084 | 8648 | 16073 | 9999 | 7 (18) |
| | | | 8632 | 16184 | . | |

**Table S3 - MinION Raw Structural Variant Prediction**

Brackets are total, number is non-LowQual. NanoSV does not try to predict type of SV (i.e. insertion, deletion etc).

| | Sanger | | MiSeq | | | |
|---|---|---|---|---|---|---|
| **Sample** | **Start** | **End** | **Start** | **End** | **QUAL** | **Total SVs** |
| N1 | - | - | - | - | - | 0 (186) |
| N3 | - | - | - | - | - | 0 (266) |
| N4 | - | - | - | - | - | 0 (345) |
| S1 | 9525 | 14387 | 9524 | 14417 | 321368.52 | 3 (300) |
| S2 | 8483 | 13459 | 8433 | 13446 | 379016.34 | 4 (286) |
| | | | 8482 | 13446 | 640446.71 | |
| | | | 8478 | 13475 | 202075.82 | |
| S3 | 5850 | 14423 | - | - | - | 0 (253) |
| S4 | 8483 | 13459 | 8447 | 13448 | 426293.68 | 2 (345) |
| | | | 8479 | 13453 | 487177.11 | |
| S5 | 9498 | 13734 | 9497 | 13722 | 708012.05 | 1 (326) |
| S7 | 7642 | 12984 | 7617 | 12986 | 143479.31 | 6 (406) |
| | | | 7638 | 12689 | 317501.45 | |
| | | | 7639 | 13014 | 155537.79 | |
| S9 | 12113 | 14421 | - | - | - | 1 (383) |
| S10 | 8649 | 16084 | 8645 | 16077 | 509797.85 | 6 (280) |
| | | | 8620 | 16072 | 419246.35 | |
| | | | 8644 | 16095 | 2333014.71 | |

**Table S4 - MiSeq Structural Variant Prediction**

Structural variants were detected by LUMPY(Express) and genotypes called using svtyper. VCF files were filtered to remove all predictions with a QUAL score of 0.00, total count is indicated in brackets.

### Supplementary Figures

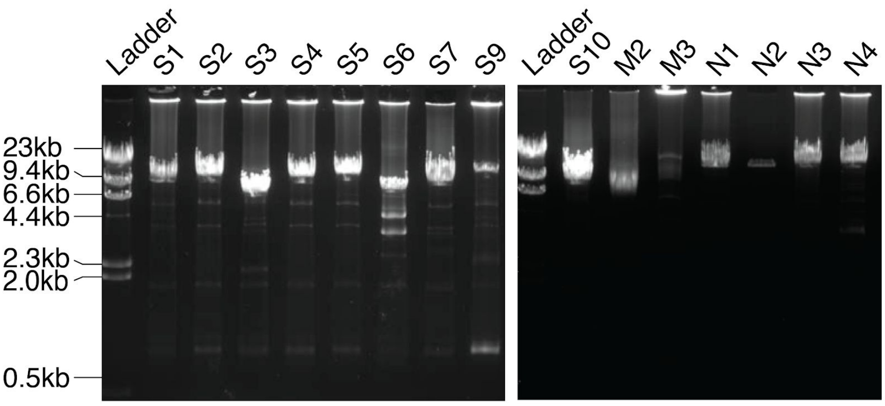


**Figure S1 - PCR of Mitochondrial DNA**

Ethidium bromide gel of the long range PCR products for each sample considered for sequencing. Samples M3 and N2 failed to produce an adequate PCR product to proceed.

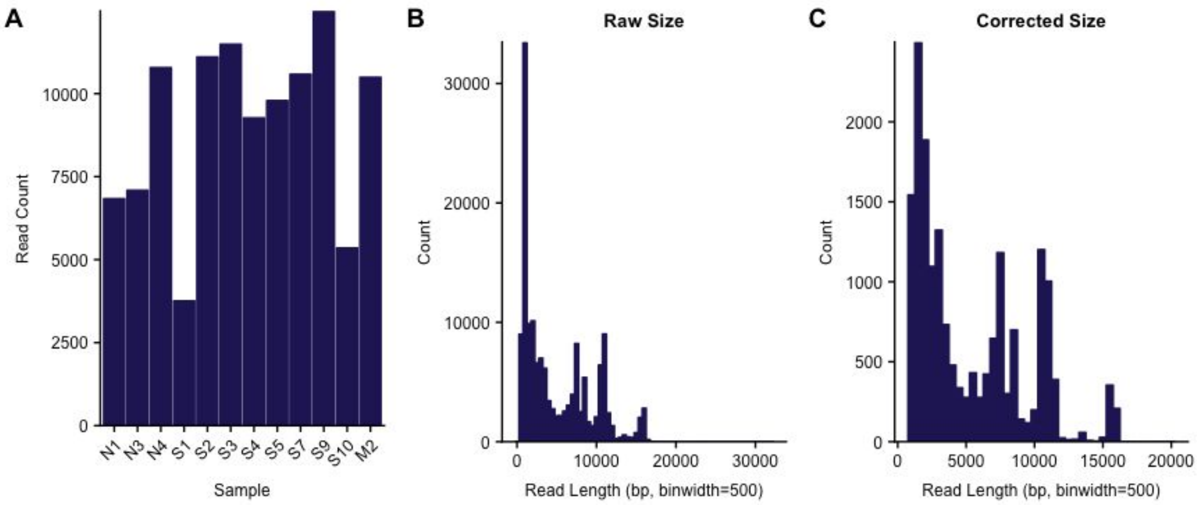


**Figure S2 - MinION Sequencing Statistics**

**A**. Number of reads for each sample after demultiplexing with poretools. **B.** Raw read length distribution of the MinION run, median read size: 3.2kb, maximum read size: 31.7kb. **C.** canu corrected and trimmed read sizes, median: 3.6kb, maximum: 19.8kb

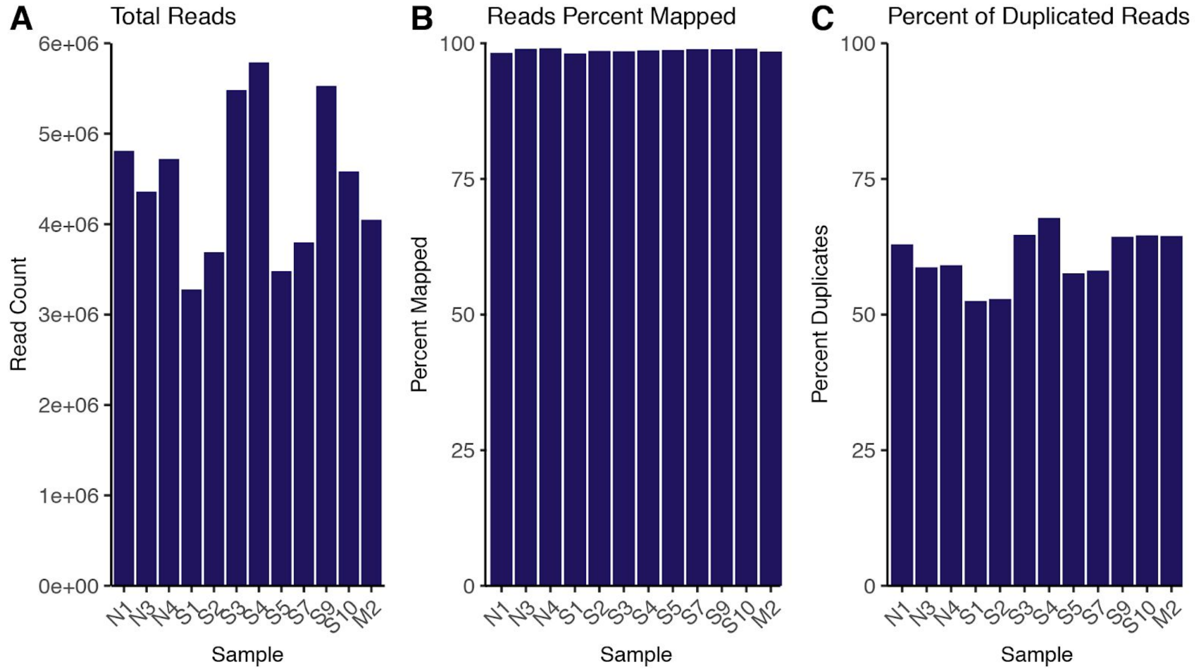


**Figure S3 - MiSeq Sequencing Statistics**

Number of total reads (**A**), percent of reads mapping (**B**) and the percentage of reads that are marked as duplicates by Picard (**C**) for MiSeq data.

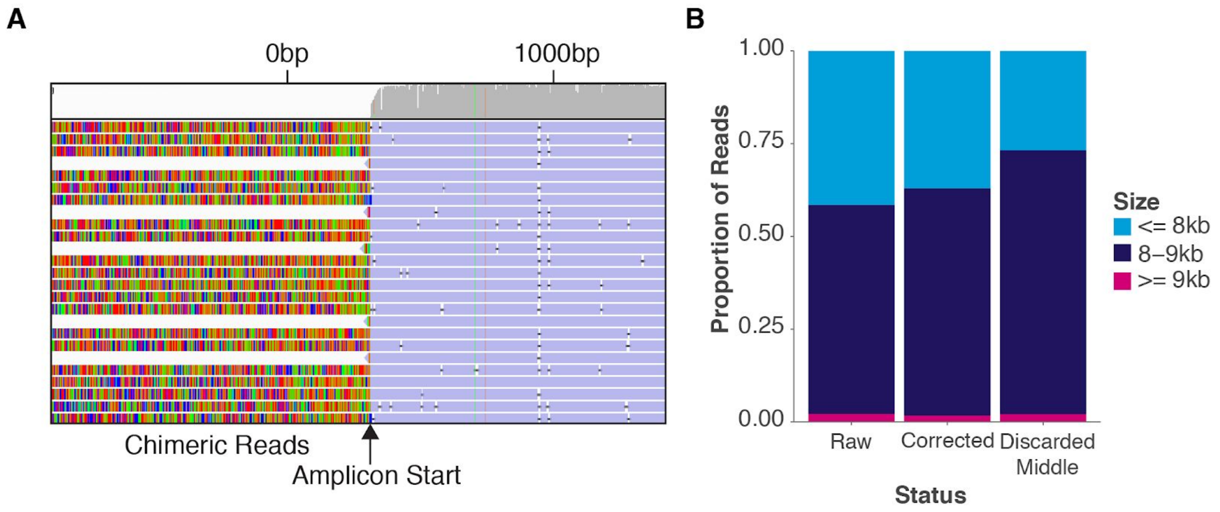


**Figure S4 - Read Lengths in Sample S10**

After the deletion is taken into account the S10 amplicon is roughly 8.5kb long. Reads longer than ~9kb are found to be chimeric reads (**A**) which contain back-to-back mitochondrial sequences, but these make up a very small percentage (2.15%) of the total. Correction reduces the amount of reads as it finds a consensus and reduces the percentage (1.7%). Using more aggressive demultiplexing with poretools which discards potentially chimeric reads that contain adapters in the middle of the sequence does not reduce the proportion of large reads (2.12%). Chimeric reads can result in bleeding of samples and so reduction of these would be advantageous in a clinical setting, in the case examined, however, all three mtDNA sequences that made up the chimera contained the same, S10, deletion.

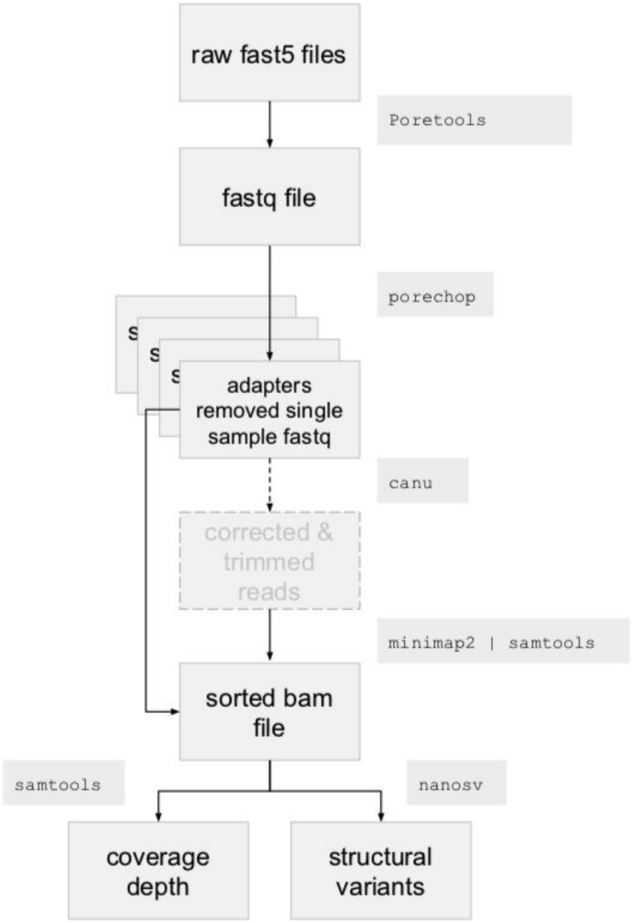


**Figure S5 - MinION SV analysis pipeline**

Flowchart depicting the analysis steps to process MinION data for mitochondrial amplicons. Briefly, a single fastq files was generated from fast5 files with poretools, porechop was used to remove adapters and demultiplex into individual sample fastq files, Optionally canu correction and trimming was performed. minimap2 was used to map reads to the rCRS reference genome, with reads sorted by samtools. Coverage depth was calculated with samtools (-Q 20), and structural variants called with NanoSV. For small variant detection a pileup file was created from the final bam file using samtools, we then used varscan to call small variants. The resulting variants were annotated with vcfanno and snpeff.

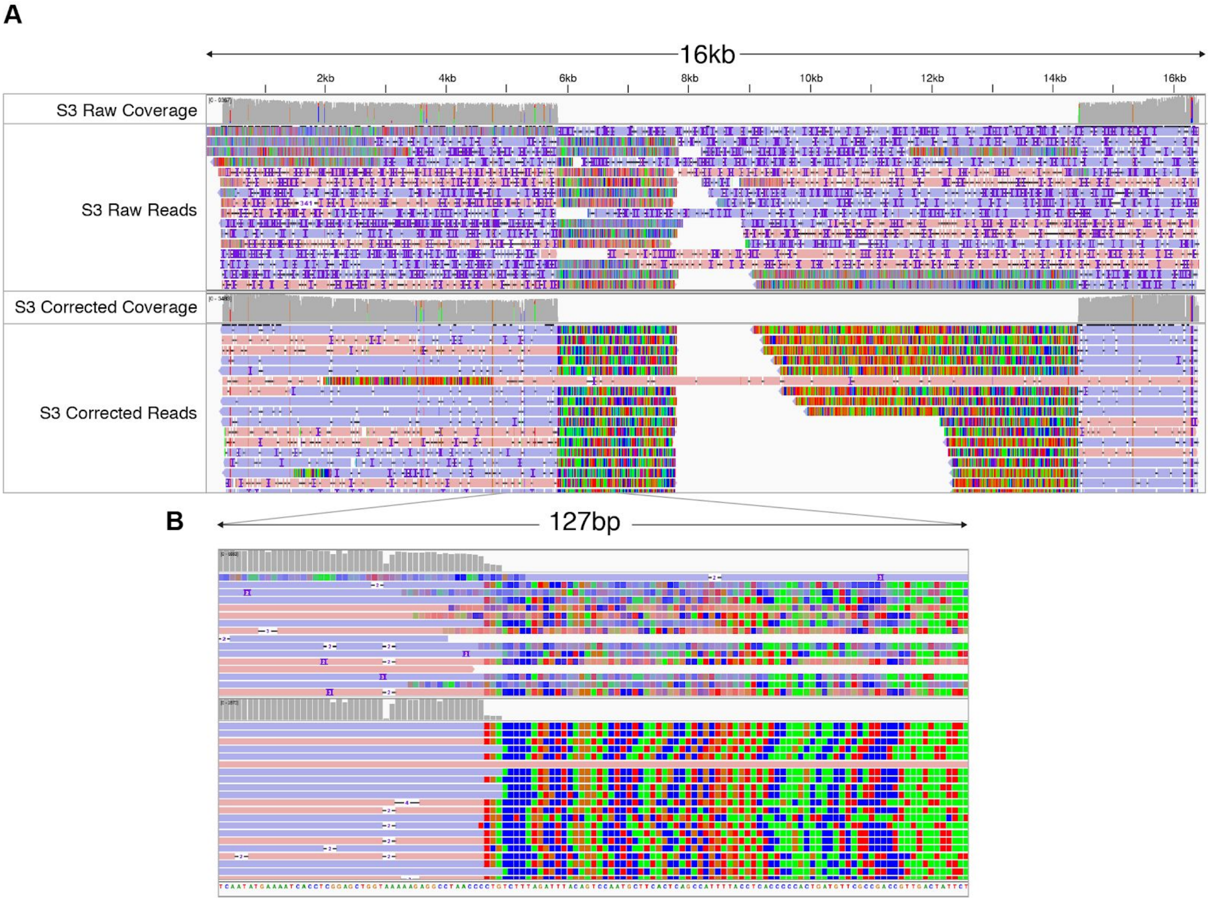


**Figure S6 - Canu Correction of MinION Reads**

Sample S3 before (top panels) and after correction and trimming with canu (bottom panels). **A.** The full 16kb amplicon in sample S3, showing high error rates and misaligned breakpoints before canu correction and trimming (top panel). After correction and trimming (bottom panel) the number of random errors are qualitatively reduced, and the position of the breakpoint is now well defined. **B.** Zoom in of a 127bp region of the amplicon showing in detail the improvement in the resolution of the breakpoint with correction and trimming (bottom panel).

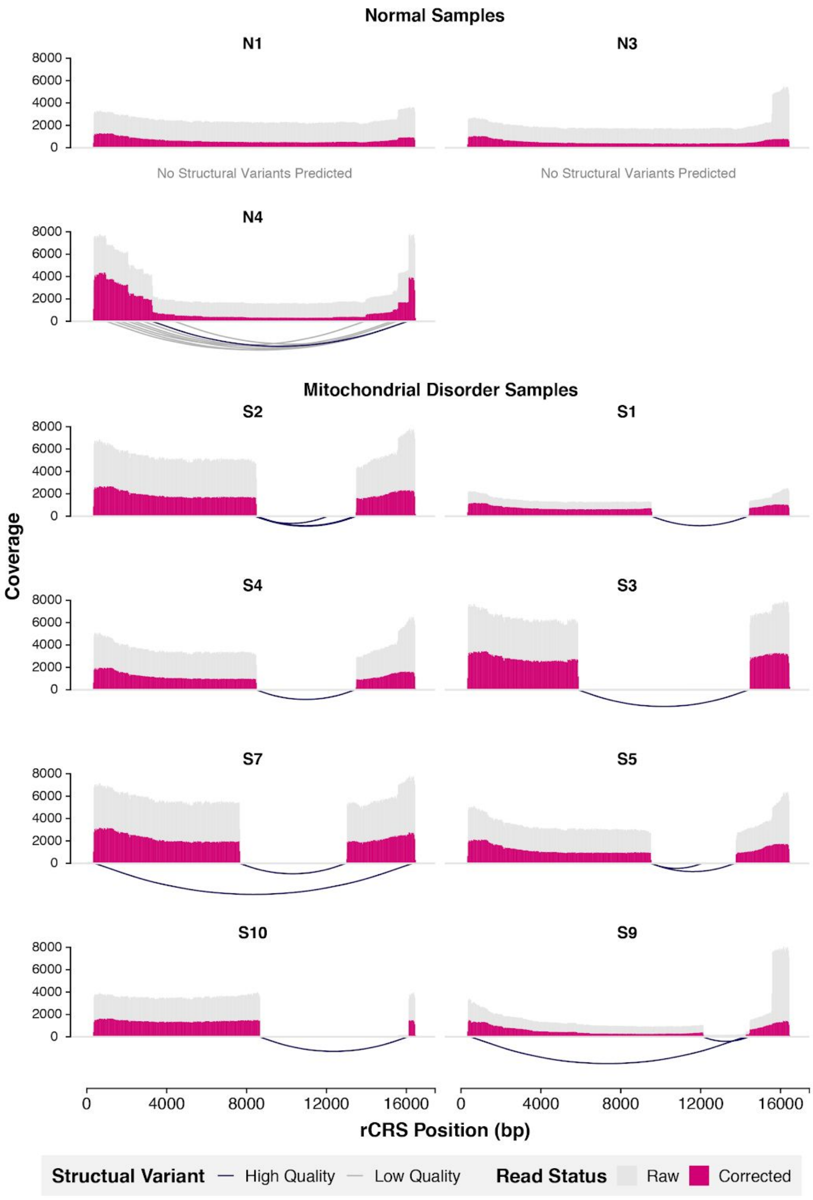


**Figure S7 - MinION Coverage and High Quality SV Predictions**

Coverage along the mitochondrial genome using both raw (grey) and canu corrected & trimmed reads (pink). Deletions are clearly represented by defined drops in coverage in all patients with mitochondrial deletions (S*). Coverage artefacts caused by short reads are visible at the starts and ends of amplicons in both raw and corrected reads, however this phenomenon is less pronounced after correction. SVs caused by artifacts at the start and end (< 320 and > 16400) of the amplicon were filtered out.

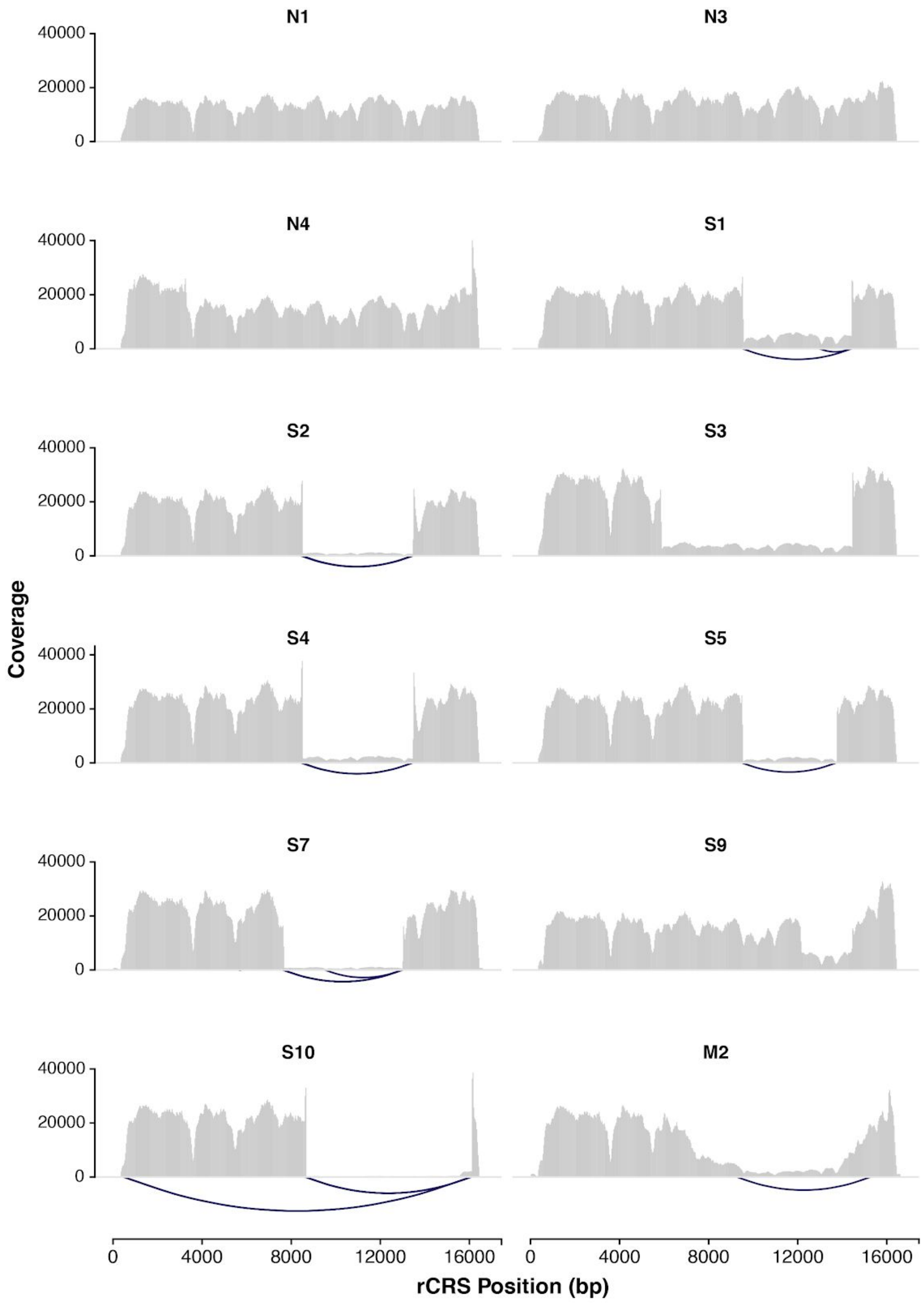


**Figure S8 - MiSeq Coverage and High Quality SV Predictions**

MiSeq coverage and high quality structural variant predictions. Compared to MinION data presented in Figure X, coverage uniformity is reduced. SVs resulting from artifacts at the start and end (< 320 and > 16400) of the amplicon were not plotted. Only SVs with a QUAL > 0.0 are displayed.

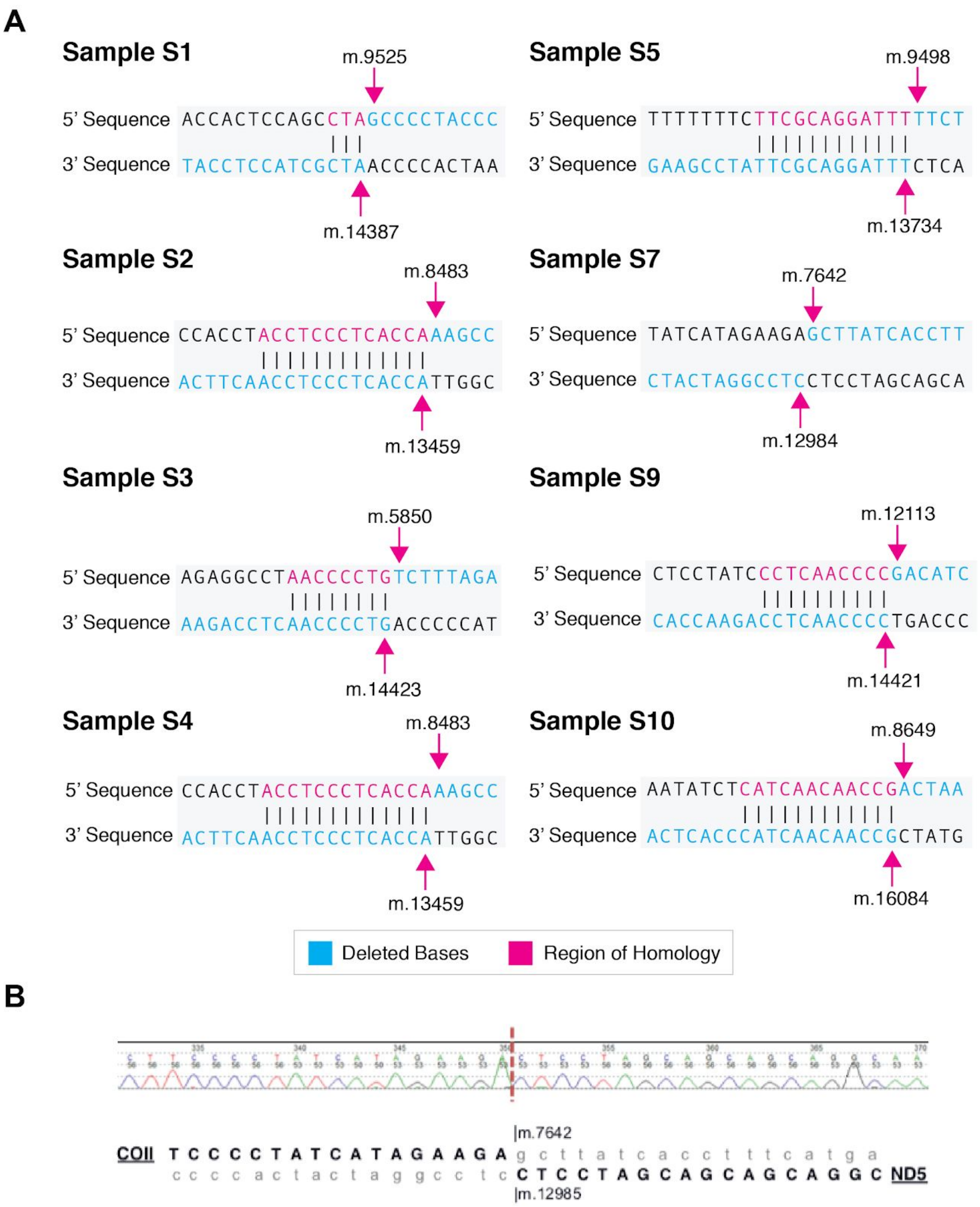


#### Figure S9 - Sequence Homology Around Deletion Breakpoints

**A.** Sequence homology around deletion breakpoints was examined manually. Sequence homology is present at the breakpoints in 7 out of the 8 single deletion samples. Sample S7 has no sequence homology. Light blue bases are those that are deleted. In accordance with variant nomenclature guidelines[1], all positions are given as if the 5' breakpoint contributes the maximum number of homologous bases to the event and the 3’ contributes none. **B.** Sequence trace for sample S7 shows the start of the deletion within COII (at position m.7642) and the end of the deletion within ND5 (at position m.12985). Lower case letters in grey indicate deleted bases.

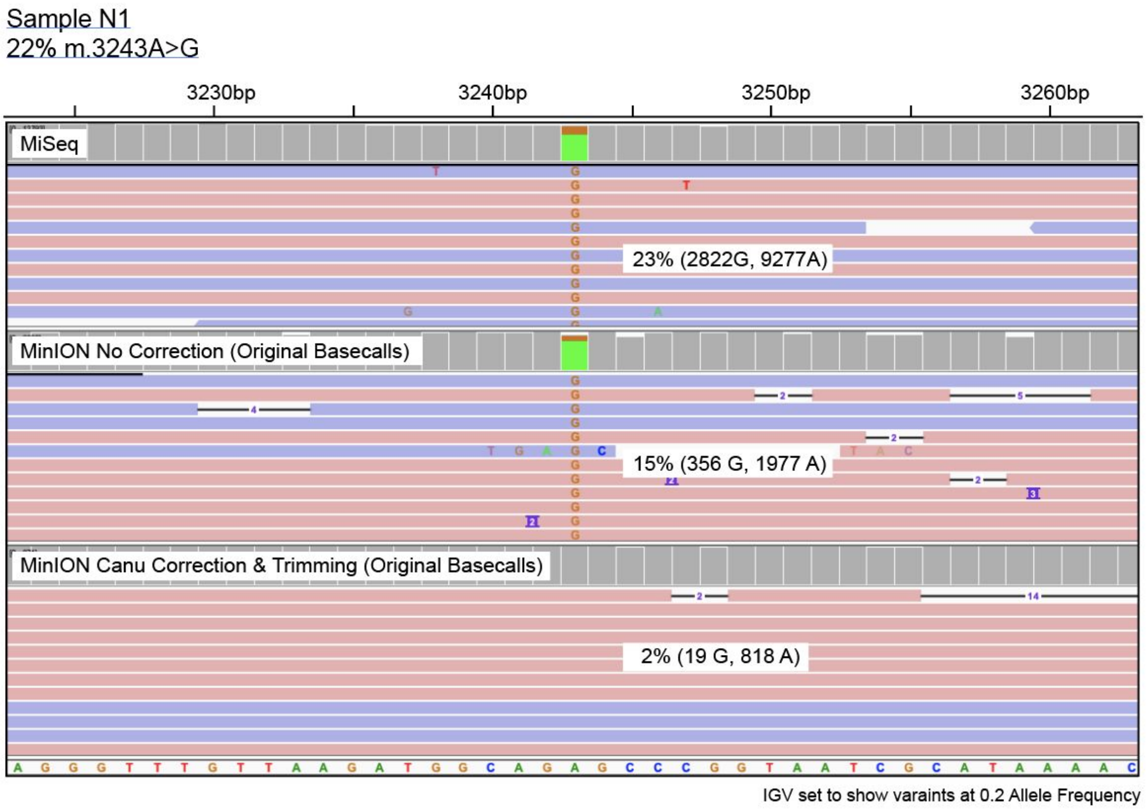


**Figure S10 - Correction causes loss of known SNV**

During the correction of reads a consensus is formed of overlapping reads, low frequency events are therefore lost as they appear like sequencing errors and are "smoothed" out in formation of the consensus. Correction on MinION reads with canu causes the reduction in the allele frequency of m.3243A>G in sample N1 of 13% resulting in only 19 reads supporting the variant at this position. This low frequency causes loss of this variant from calls.

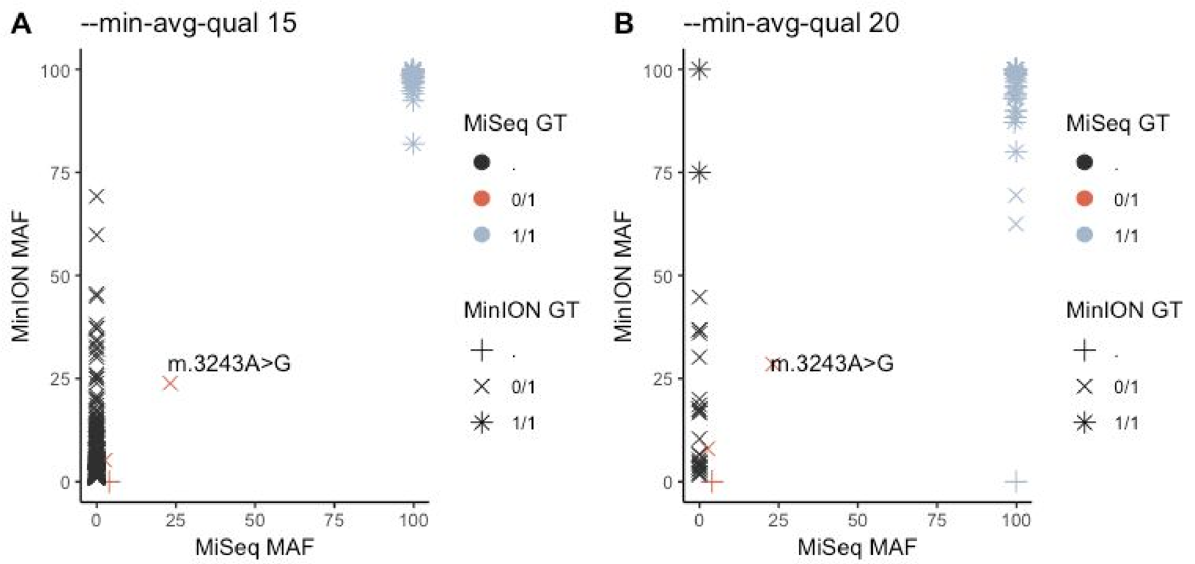


C

| Class | Total.SNPs |
|---|---|
| MiSeq only | 1 |
| MinION only | 368 |
| Both | 33 |

D

| Class | Total.SNPs |
|---|---|
| MiSeq only | 2 |
| MinION only | 27 |
| Both | 32 |

**Figure S11 - Known variants are called by VarScan using MinION data**

Calling variants with VarScan in sample N1 for both MiSeq and MinION data shows reasonable concordance. Plots A & B show the minor allele frequency of variants present in both technologies and those only present in one. Using a minimum average quality of alternate bases of 15 (A, C) all but 1 variant present in the MiSeq is called as present in the MinION data, however a high number of additional variants are present in the MinION data aone (368). Applying a more stringent base quality minimum of 20 reduced false positives to 27, but with the gain of a false negative. The known variant m.3243A>G (22%) is retained in both samples at an expected allele frequency.

(Data here is from albacore called bases)

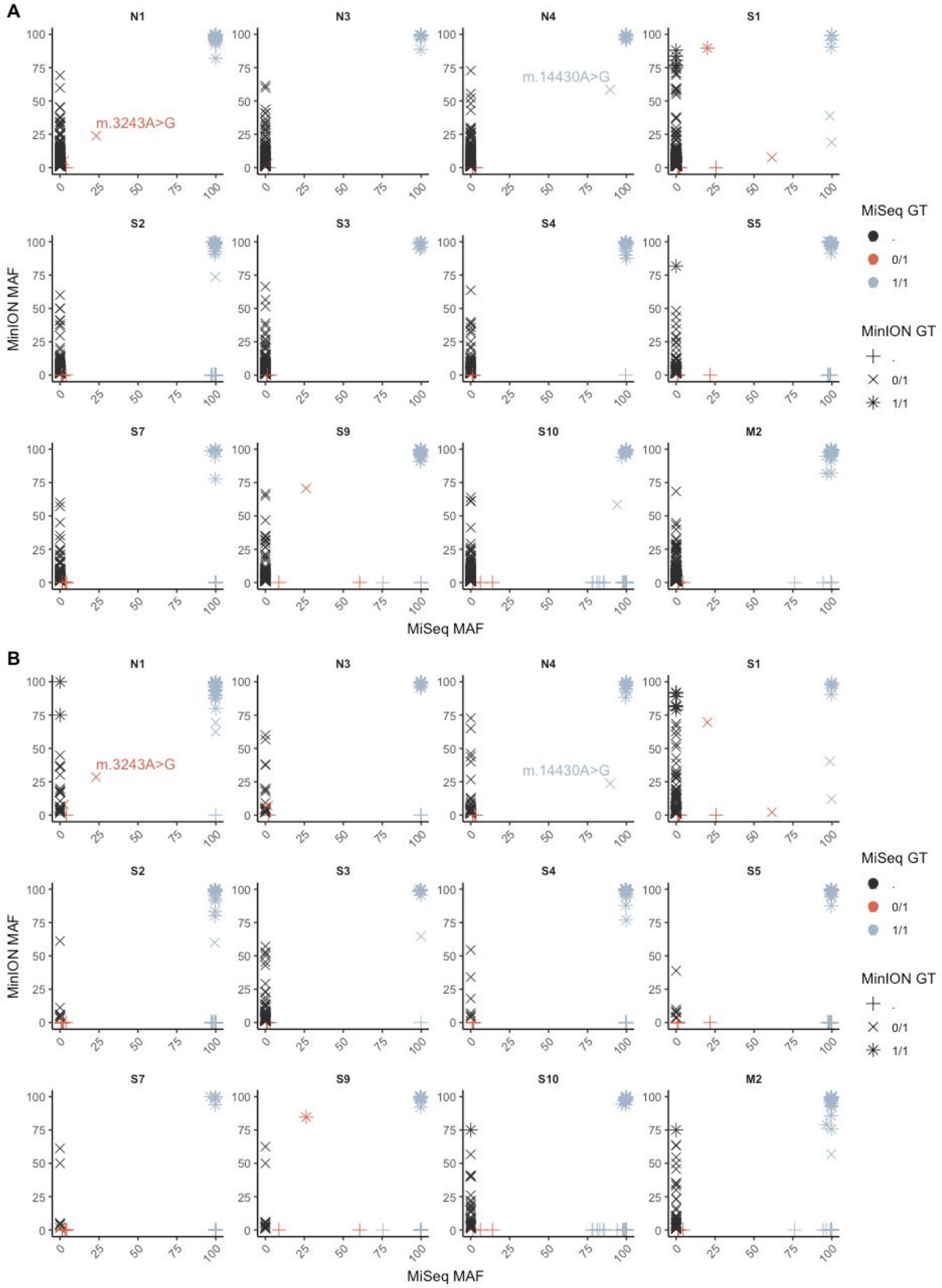


**Figure S12 - Variant Calling Comparison for All Samples**

**A.** Using minimum average base quality of 15. **B.** Using minimum mapping quality of 20

Some of the missing variants in MinION are found in the deleted region in MiSeq or at the starts and ends of the genome where we have lower coverage in MinION., For example sample S10 has 20 1/1 calls missing in MinION in the min-ave-quality 20 dataset. 13 of these are in the deletion and therefore have low coverage in MinION, whereas 6 are at the starts or ends of the sequence and have low coverage. The remaining call is co-located with a deletion causing issues for the variant caller. 3 0/1 calls are missing, and these are all located in the deleted region. There are no other false negatives or mismatched genotypes in this sample leaving 43 calls which are not found in MiSeq data.
